# Accurate hybrid plasmids assembly with HyPlAs

**DOI:** 10.1101/2025.08.21.671630

**Authors:** Fatih Karaoglanoglu, Kay C. Wiese, Cedric Chauve

**Affiliations:** School of Computing Science, Simon Fraser University, 8888 University Drive, Burnaby, V5A 1S6, BC, Canada; Department of Mathematics, Simon Fraser University 8888 University Drive, Burnaby, V5A 1S6, BC, Canada

**Keywords:** Plasmids, hybrid short- and long-reads sequencing, genome assembly

## Abstract

**Summary:** The increasing availability of hybrid sequencing datasets comprising both short and long reads, is radically transforming microbial genomics, with the prospect of obtaining routinely near-complete bacterial genome assemblies. However, the complete assemblies of mobile genetic elements, especially plasmids, still remain challenging. We introduce HyPlAs, an assembly pipeline specifically designed to assemble plasmids from hybrid bacterial sequencing datasets. HyPlAs main novelty is to incorporate the use of a prior classification of short-read contigs as chromosomal or plasmidic. We evaluate HyPlAs on a large set of bacterial samples and demonstrate that it outperforms its competitor Plassembler.

**Availability:** HyPlAs is freely available at https://github.com/cchauve/HyPlAs.

## Introduction

Plasmids are extrachromosomal DNA sequences that can carry antibiotic-resistant genes and are a major vector of the spread of antimicrobial resistance (AMR) [Castañeda-Barba et al., 2024, Vrancianu et al., 2020], a major public health concern. This motivates the development of efficient and accurate bioinformatics methods that can detect plasmids present in the sequencing data generated by epidemiological surveillance protocols [Sobkowiak et al., 2025].

The detection of plasmids from short-read sequencing data is an active research area, focusing either on identifying contigs originating from plasmids (plasmid contigs classification) [Sielemann et al., 2023, Schwengers et al., 2020, Tang et al., 2023, Andreopoulos et al., 2021, Arredondo-Alonso et al., 2018, van der Graaf-van Bloois et al., 2021], or on clustering contigs into groups likely originating from the same plasmid (plasmid binning) [Rozov et al., 2016, Robertson and Nash, 2018, Paganini et al., 2024] or on assembling plasmid sequences [Wick et al., 2017b, Antipov et al., 2016].

Hybrid (short-read and long-read) data are increasingly used in an epidemiological setting (e.g. [Foster-Nyarko et al., 2023]). Despite the potential of hybrid sequencing data to generate near-complete assemblies [Wick et al., 2023], recovering fully assembled plasmids present in bacterial isolates remains challenging in some aspects [Johnson et al., 2023]. However very few methods are available to process hybrid data for bacterial samples with a specific focus on plasmids. Unicycler [Wick et al., 2017b,a] has long been the main tool used to recover plasmids from hybrid data, as a by-product of full bacterial genomes assembly. The recently introduced tool Plassembler is the only method we are aware of with a specific focus on plasmids recovery [Bouras et al., 2023]. Plassembler proceeds in thee main stages: (1) it first assembles long-read data to identify the chromosome and putative plasmids, (2) it maps short reads and long reads to this assembly to select reads possibly originating from plasmids (*plasmidic reads*), (3) it assembles from scratch the selected plasmidic reads using Unicycler, resulting in plasmid contigs.

We introduce HyPlAs, a method aimed at assembling plasmids from hybrid data. HyPlAs is based on three guiding principles. First, if one can extract from a hybrid dataset all plasmidic long reads, provided that sequencing coverage is high enough, then plasmids can be obtained by refining the short-read assembly using these plasmidic long reads and extracting circular contigs. Second, we hypothesize that, when detecting putative plasmidic long reads to refine the short-read assembly, a high recall is more important than precision: if most plasmidic long reads have been selected they will result in circular contigs and chromosomal long reads incorrectly selected as plasmidic will result in linear contigs that can be ignored. Third, short-read plasmid contigs classification can be leveraged to identify plasmidic long reads, an avenue that has not been explored to date: long reads that contain plasmidic short-read contigs are likely to be plasmidic long reads. We evaluate the performance of HyPlAs on hundreds of hybrid datasets for bacterial samples and show that it does outperform Plassembler.

## Methods

HyPlAs is a pipeline combinig existing tools and specific Python scripts and C++ programs. HyPlAs is composed of the following steps (illustrated in Fig. 1): (1) Reads preprocessing; (2) Short reads assembly using Unicycler [Wick et al., 2017b]; (3) Detection of putative plasmidic long reads, done in four stages: (3.a) the plasmid contigs classification tool Platon [Schwengers et al., 2020] is used to detect plasmidic short-read contigs, (3.b) long reads are mapped to the assembly graph using minigraph [Li et al., 2020], (3.c) long-read mapping to short-read contigs and Platon results are used to select an initial set of putative plasmidic long reads, (3.d) the set of putative plasmidic long reads is augmented by iteratively detecting overlapping long reads; (4) Refinement of the short-read assembly graph generated in step 2 using the plasmidic long reads selected during step 3. We describe in more detail the various steps of HyPlAs below.

**Fig. 1.**
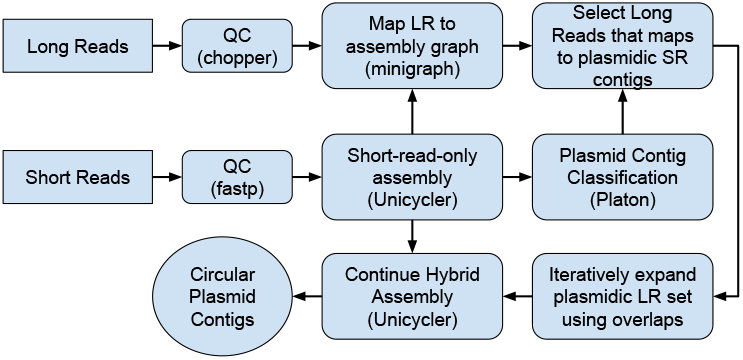
HyPlAs pipeline architecture

### Reads preprocessing

Similarly to Plassembler, reads are filtered and trimmed using fastp [Chen et al., 2018] for short reads and chopper [De Coster and Rademakers, 2023] for long reads; the parameters values used are the same than in Plassembler.

### Short-read assembly

We assemble the short-read data using Unicycler [Wick et al., 2017b]. While other bacterial genome assemblers that generate an assembly graph, such as SKESA [Souvorov et al., 2018] or SPAdes [Bankevich et al., 2012], could be used, we chose Unicycler as the default as it is also used for the final hybrid assembly step in the pipeline.

### Plasmidic long reads detection

The selection of a subset of long reads that include the long reads that are assumed to have been sequenced from actual plasmids (termed plasmidic long reads) is the heart of HyPlAs.

#### Short-read contig classification

In order to detect plasmidic long reads, HyPlAs first labels short-read contigs as either *plasmidic, chromosomal* or *ambiguous*. To do so, HyPlAs relies on using the plasmid contigs classification tool Platon [Schwengers et al., 2020]; we chose Platon over other classification tools as it is based on an interpretable decision tree, unlike most other classification tools that rely on machine learning.

Platon contigs classification procedure can be summarized as follows. The replicon distribution score (RDS), based on the comparison of contigs against a database of plasmid genes, is the main parameter of the method, where a high RDS indicates a contig likely to be plasmidic, a low RDS a contig likely to be chromosomal and contigs with an intermediate RDS are classified as plasmidic if they exhibit at least one of a series of features indicating a likely plasmid origin (such as the presence of a mobility protein gene or being circular). We classify contigs based on the results of Platon as follows: (1) a contig classified as plasmidic by Platon is labeled as plasmidic, (2) a contig with a low RDS score is labeled as chromosomal, and (3) a contig with an intermediate RDS score and exhibiting none of the plasmid features considered by Platon is labeled as ambiguous.

#### Putative plasmidic long reads selection

Given a short-read draft assembly, a classification of its contigs as either plasmidic, chromosomal or ambiguous, and long-read data, we start by mapping the long reads to the assembly graph using minigraph [Li et al., 2020]. For each long read, minigraph reports either “nothing” if there is no mapping, “single alignment” if the read can be mapped to the graph by using the edges in the assembly graph, or “multiple alignments” where the read needs to be broken down due to lack of edges that could connect partial alignments against the contigs.

Based on this mapping, we label a long read as *plasmidic* if its mapping to the assembly graph does not include any chromosomal short-read contig and includes at least one plasmidic short-read contig. Long reads with no mapping are labelled as *unmapped*, long reads with a mapping that contains no chromosomal or plasmidic short-read contig are labeled as *unknown* and long reads with mappings to both plasmidic and chromosomal contigs are labelled as *ambiguous*; last, long reads whose mapping contains at least one chromosomal contig and no plasmidic contigs are labeled as *chromosomal*. We select the subset of long reads labeled as plasmidic as the initial set of putatively plasmidic long reads.

However, we observed that while this procedure recovers a large part of the true plasmidic long reads, some of the larger plasmids were not fully covered by the selected long reads, which resulted in their incomplete assembly. To address this issue, HyPlAs augments the initial set of putative plasmidic long reads by detecting additional long reads that overlap with them. Unmapped, unknown and ambiguous, long reads are mapped to the set of already identified plasmidic long reads using minimap2 [Li, 2018] and any ambiguous long read with such a mapping is added to the putative plasmidic long-read set. This augmentation step can be skipped or iterated any number of times to trade precision with recall. In our experiments, we denote HyPlAs-0, HyPlAs-1 and HyPlAs-2 the method HyPlAs without the augmentation step, with one augmentation iteration and two augmentation iterations respectively.

### Final hybrid assembly

HyPlAs concludes by using Unicycler [Wick et al., 2017b] in hybrid mode, with the selected putative plasmidic long reads and the initial short-read assembly graph, so the initial short-read assembly is refined using the set of putative plasmidic long reads. HyPlAs then reports the circular contigs in the Unicycler output as predicted plasmids.

## Results

To compare HyPlAs against Plassembler, we applied both tools to a set of hundreds of bacterial sequencing datasets for which complete assemblies, short-read data and long-read data were publicly available. We evaluated the methods on two criteria: accuracy of the selection of putative plasmidic long reads and accuracy of the reported assembled plasmids.

### Data

We focused on the ESKAPEE pathogens group [Miller and Arias, 2024], that consists of 7 bacterial species of high interest in terms of pathogenicity and antimicrobial resistance.

We used the NCBI Genome utility [Kitts et al., 2016] to collect data for all ESKAPEE samples satisfying the following criteria: (1) complete assembly, (2) released between 2012 and 2024, (3) short-read and long-read Oxford Nanopore sequencing datasets publicly available in the NCBI Sequence Read Archive (SRA). In order to keep the dataset consistent, we focused on the subset of 590 samples with at least one annotated plasmid and assembled using Unicycler. Finally, we removed 30 samples from the analysis due to Plassembler terminating with an error. This left 560 samples with 1481 plasmids.

We labeled the 1481 ground truth plasmids as *small* if their length was below 20000bp and *large* otherwise.

We processed these samples with HyPlAs-0, HyPlAs-1 and HyPlAs-2, and Plassembler (with default parameters, but in both versions using the long-read assemblers Flye [Kolmogorov et al., 2019] and Raven [Vaser and Šikić, 2021]).

### Plasmid recovery evaluation

We first compare the ability of HyPlAs and Plassembler to recover true plasmids. To do so we define a metric allowing us to compare the ground truth plasmids and the plasmids predicted by HyPlAs and Plassembler. We compute the Jaccard similarity between all plasmids in the ground truth and the plasmids reported by each tool. Then, starting from the highest similarity pair in decreasing order, we matched two plasmids (a ground truth plasmid and a predicted plasmid) if they have at least 75% similarity and have not already been matched with another true or predicted plasmid. We then define 3 values: *G* ← *# of ground truth plasmids, P* ← *# of predicted plasmids*, and *M* ← *# of ground truth matched Plasmids*. Using these we calculate the precision as 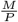, the recall as 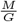 and the F1-score as 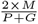.

Table 1 shows the performance of the tools on the 560 samples and 1481 plasmids. We can see that HyPlAs outperforms Plassembler slightly in terms of recall and F1-score and predicts 28 additional true plasmids with only 2 additional false positives (HyPlAs-2). Plassembler performs better when run with Flye, thus, in the remaining analysis, we only consider the Plassembler results obtained with Flye.

**Table 1.** Plasmid prediction accuracy statistics. TP: number of true positive predicted plasmids (|*M* |). FP: number of false positive predicted plasmids (|*P* − *M* |). FN: number of false negatives (|*G* − *M* |).

|  | Precision | Recall | F1-score | TP | FP | FN |
| --- | --- | --- | --- | --- | --- | --- |
| HyPIAs-0 | 0.8987 | 0.6111 | 0.7275 | 909 | 98 | 572 |
| HyPIAs-1 | **0.9093** | 0.8805 | 0.8947 | 1311 | **123** | 170 |
| HyPIAs-2 | 0.9066 | **0.8845** | **0.8954** | **1318** | 131 | **163** |
| Plassembler-Raven | 0.8934 | 0.8258 | 0.8582 | 1222 | 141 | 253 |
| Plassembler-Flye | 0.9056 | 0.8677 | 0.8862 | 1290 | 129 | 191 |

To better understand the limitations of both methods, we separated true plasmids into four sets: recovered by both methods (*HP*), only missed by HyPlAs (*H*^*′*^), only missed by Plassembler (*P* ^*′*^), and missed by both methods (*HP* ^*′*^). These sets contain 1228, 57, 82 and 114 plasmids respectively. We analyzed these sets on two plasmid attributes: length and similarity with the chromosome.

Of the 641 (resp. 840) small (resp. large) plasmids, 556 (resp. 672) were recovered by both methods, 22 (resp. 35) were missed by only HyPlAs, 21 (resp. 61) were missed only by Plassembler and 42 (resp. 72) were missed by both. Thus, we did not observe a significant difference between the performance of both methods based on plasmid length.

Next, we analyzed the effect of plasmid/chromosome sequence similarity using Jaccard containment [Koslicki and Zabeti, 2019]. Using this metric, we observed (Figure 2) that sequences of the plasmids missed by Plassembler have higher similarity with the chromosome(s). The mean containment values for these plasmids sets were 0.030 (resp. 0.049, 0.082, 0.049) for plasmids in *HP* (resp. *H*^*′*^, *P* ^*′*^, *HP* ^*′*^). The difference between the values for *H*^*′*^ and *P* ^*′*^ is significant according to a Wilcoxon-Mann-Whitney test (*p* = 0.001). This result shows that Plassembler seems more prone to miss plasmids with high chromosome similarity compared to HyPlAs. This can be observed for both small and large plasmids, but it is more pronounced for small plasmids.

**Fig. 2.**
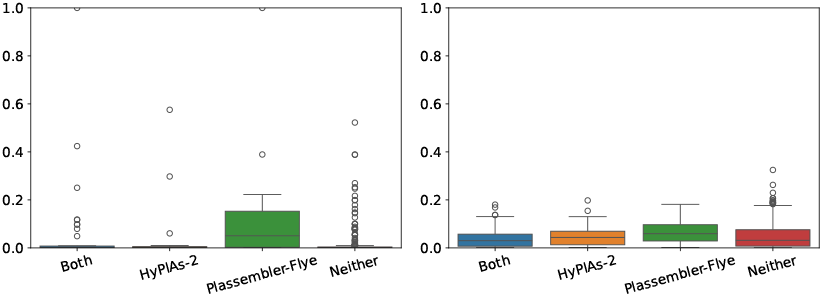
Bar-plot of the sequence containment ratio distribution of (Left) small plasmids and (Right) large plasmids missed by neither of the methods, by Plassembler, by HyPlAs-2, or by both.

### Plasmidic long reads selection evaluation

The second aspect we evaluated is the ability of HyPlAs and Plassembler to identify plasmidic long reads.

We first defined true plasmidic long reads for each sample by mapping all long reads to the ground truth plasmid sequences and selecting reads with at least 90% mapping ratio (number of mapped bases divided by the read length), then compared this true plasmidic read-set to the long reads selected by Plassembler and HyPlAs as plasmidic. Supplementary Figure 6 shows the results of this experiment. We observe that generally Plassembler finds more plasmidic long reads than HyPlAs. To assess how well the selected plasmidic long reads were actually covering the plasmids present in the samples, we calculated the sequence coverage of the ground truth plasmids by the plasmidic long reads mapped to them as described above. Figure 3 shows that HyPlAs is able to recover plasmidic long reads that result in an improved coverage of the true plasmids.

**Fig. 3.**
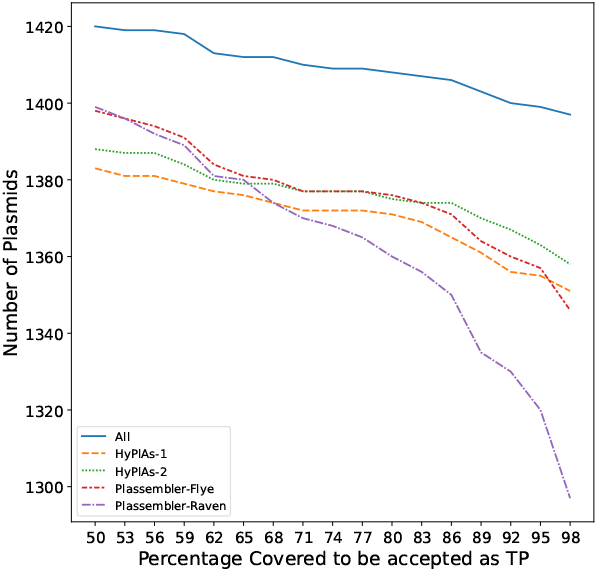
Number of plasmids covered over at least *x*% of their length (*x*-axis) by selected plasmidic long reads. At *x* = 90%, Plassembler-Raven recovers 1369 plasmids out of 1442, Plassembler-Flye recovers 1401 plasmids, HyPlAs-0 1397, HyPlAs-1 1422 and HyPlAs-2 1425.

**Fig. 4.**
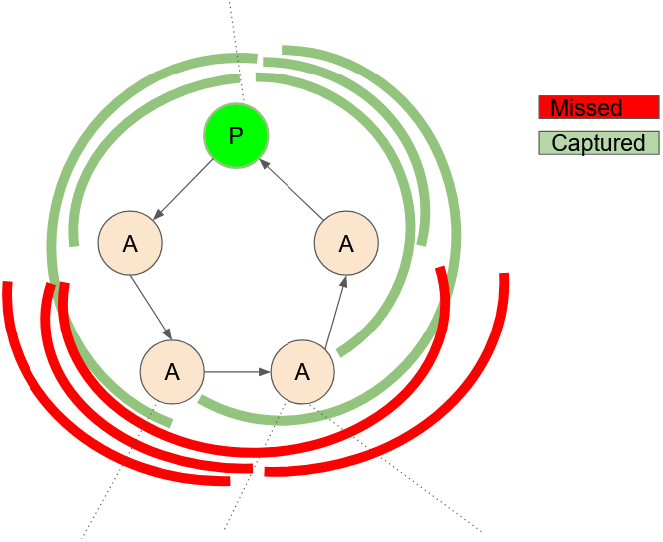
Illustration of long reads that may be missed during the initial plasmidic long read selection step. Given plasmid sequences are often composed of cycles of contigs in the short-read assembly graph and they are predicted as either plasmid (P) or ambiguous (A): If there are long stretches of ambiguous contigs, some of the long reads originating from this plasmid may be missed during the read selection if they do not overlap with any contig classified as plasmidic(Colored red).

## Discussion

HyPlAs is a novel plasmid assembler that follows the same general principles than Plassembler but introduces novelties, notably the use of a short-read contigs classification tool to help detect putative plasmidic long reads, and the idea to refine the short-read assembly graph with predicted plasmidic long reads to assemble plasmids.

Our experiments show that, when aiming to detect plasmidic long reads, to be used to refine the short-read assembly to identify plasmids, allowing a significant loss of precision to increase the recall seems to be a valid strategy. Indeed, it results in an increased coverage of plasmids by long reads at the expense of including chromosomal long reads that do not form spurious cycles in the assembly graph. Nevertheless, we believe that more accurate methods to detect plasmidic long reads, a problem that has not been considered so far to the best of our knowledge, would have a positive impact.

Our experiments also show that Plassembler is more likely to miss plasmids with higher chromosome similarity. We suspect this is due to Plassembler plasmidic read selection process, where it filters out true plasmidic read with chromosome mapping. HyPlAs, on the other hand, selects the reads that map to plasmid contigs in the assembly graph and allows mapping to the ambiguous contigs. This approach allows HyPlAs to recover plasmid reads with chromosome similarity provided they do overlap with high-confidence plasmidic reads identified through using Platon.

Although HyPlAs shows improved performance on plasmids with high chromosome similarity, several plasmids undetected by HyPlAs were identified by Plassembler. We did not find any discernible pattern among these missed plasmids.

## Accurate hybrid plasmids assembly with HyPlAs (supplementary material)

### Supplementary methods

#### Platon

By default, Platon only reports plasmid contigs, where any omitted contig is assumed to be chromosomal. Since it was optimized for accurate plasmid classification, this was not an oversight by the authors. Nevertheless, on top of the accurate plasmid classification, HyPlAs requires accurate classification of the chromosomal contigs as well. To achieve this, we implemented a slight modification of the decision procedure of Platon. Note that this change did not require modifying the code; rather, it was done by using the “–characterize” option, which, instead of making decisions, outputs all of its decision parameters per input contig. Platon plasmid classification procedure can be summarized as follows. The replicon distribution score (RDS) is the main parameter of the method where a high RDS indicates a contig is likely to be from a plasmid. Authors empirically determine two thresholds for the RDS value optimizing the plasmid classification referred to as sensitivity threshold (SNT) and specificity threshold (SPT). The sensitivity threshold is then used to exclude chromosomal contigs: if the RDS value of a contig is less than SNT, it will be considered chromosomal. Any contig with the RDS value above the SPT will be classified as plasmidic.

In summary, Platon uses the Sensitivity threshold to determine chromosomal contigs and the Specificity threshold to determine plasmid contigs. For any contig with RDS value between these thresholds, a series of checks are considered (such as circularity, mobilization or replication proteins), and a contig is labeled plasmid if any of the checks are valid and chromosome otherwise (the flowchart describing this procedure can be found in [Schwengers et al., 2020, Fig. 1]). Our procedure simply changes the final step of this decision and labels a contig as ambiguous if it has an RDS value between these two thresholds and it does not pass any of the plasmid checks.

#### Plasmidic long reads selection

We use the following Algorithm to label long reads using the alignments generated by minigraph.

##### Algorithm 1

Labeling of long reads

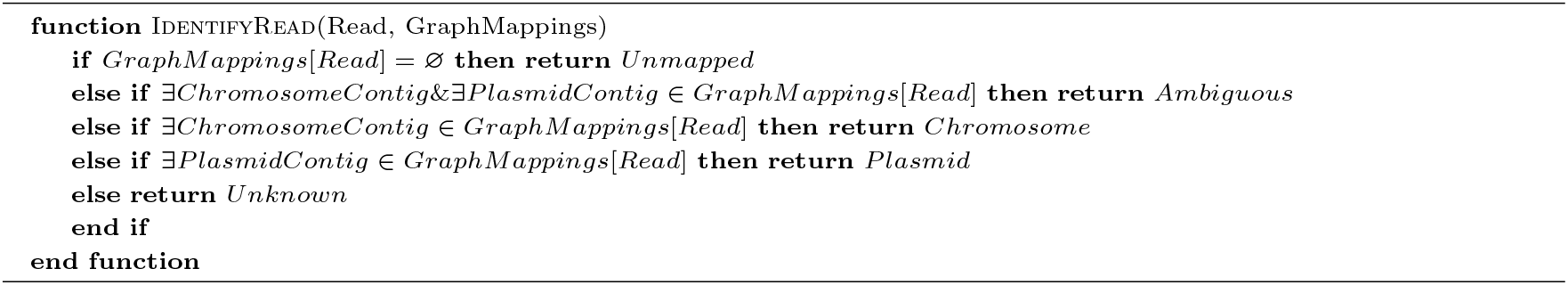

The long reads labeled as *Plasmid* and *Unknown* are then used for the final plasmid assembly.

#### Jaccard similarity and Jaccard containment index

To compute the Jaccard containment index of a plasmid *p* compared to a chromosome *c*, we first compute the minhashes of both the chromosome and the plasmid using sourmash [Pierce et al., 2019] with parameters (K=17, Scale=15). Using these minhashes, we estimated the Jaccard containment [Koslicki and Zabeti, 2019] of the plasmid sequences within their main chromosomes as 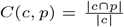 where *c* and *p* represents the *k* −*mer* sets of the chromosome and the plasmid respectively. This metric is more suitable than the Jaccard similarity, since it is not affected by size difference and extra sequences in the chromosomes.

### Supplementary results

#### Ground truth plasmid length and coverage analysis

We observed that the distribution of the length of the plasmids from the ground truth on the 560 samples is bimodal, with very few predicted plasmids of length between 10,000 and 30,000 base pairs(Figure 5). Following this, we used 20000 base pairs as a breakpoint to label plasmids as either small or large plasmid. Then, for each plasmid, we computed its relative short-read coverage (compared to the chromosome of the sample) and compared it to its length in a scatter-plot (Figure 5a). We notice that small and large plasmids follow different coverage distributions where short plasmids had many copies and large plasmids had few, which agrees whith what is known about plasmids [Dewan and Uecker, 2023].

**Fig. 5.**
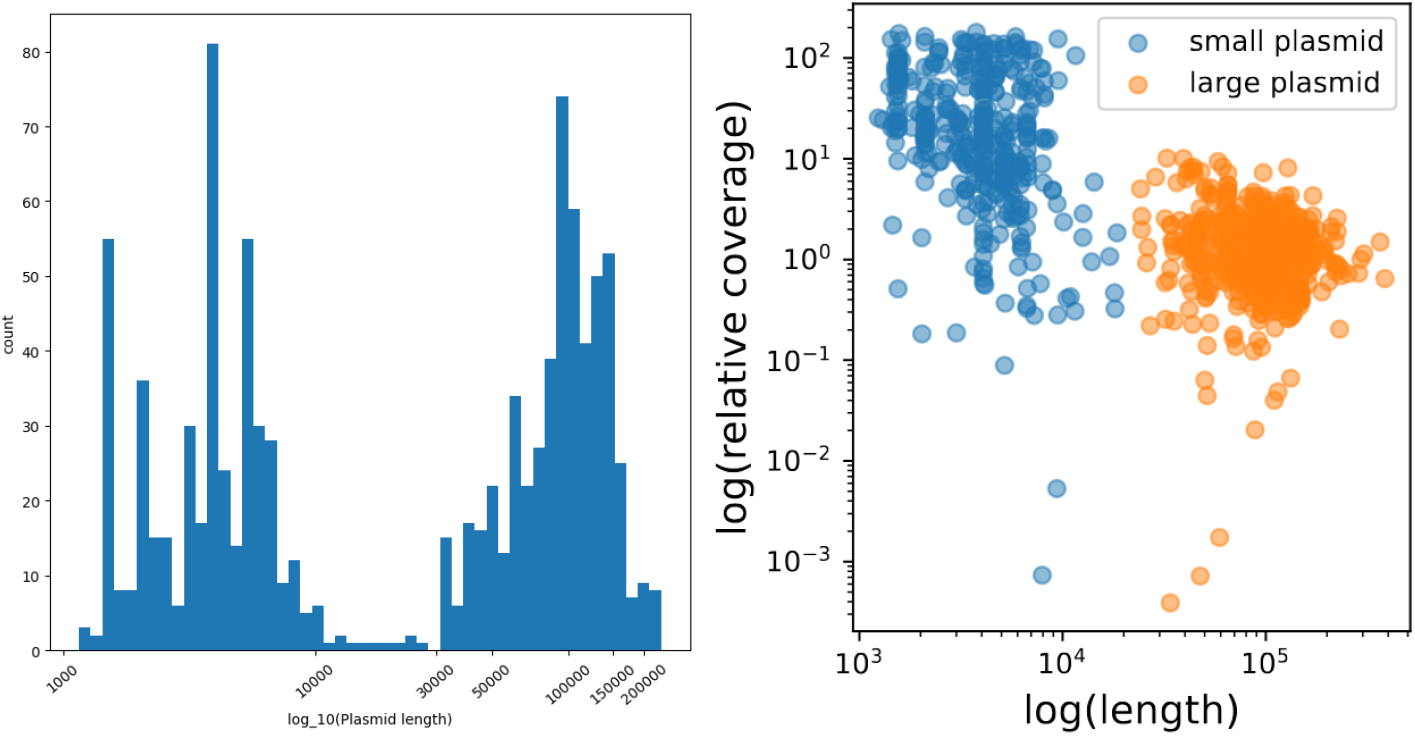
(Left) Histogram of ground truth plasmid length. (Right) Scatter plot of the normalized sequening coverage of small and large plasmids

**Fig. 6.**
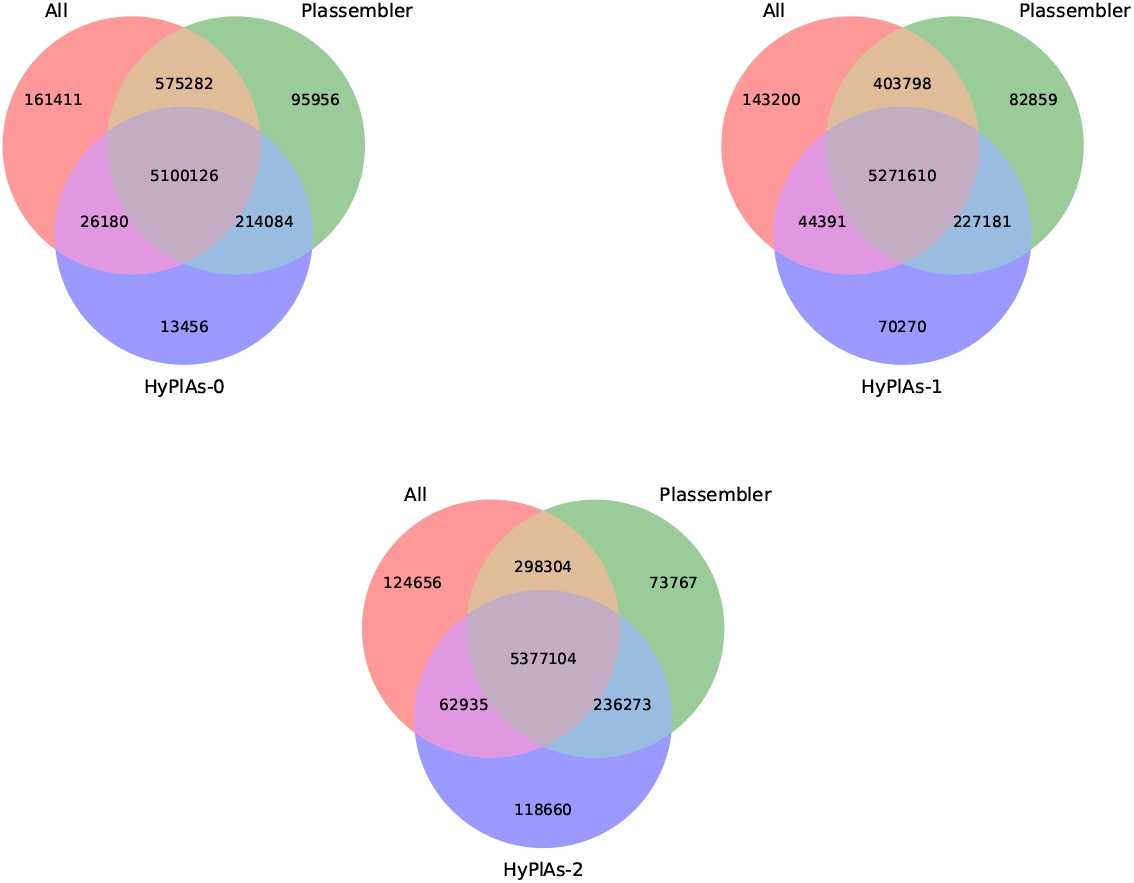
Venn diagram of HyPlAs predicted plasmidic long reads, Plassembler predicted plasmidic long reads and ground truth plasmidic long reads. (Top Left), (Top Right), (Bottom) illustrate respectively the results using HyPlAs-0, HyPlAs-1, and HyPlAs-2.

**Fig. 7.**
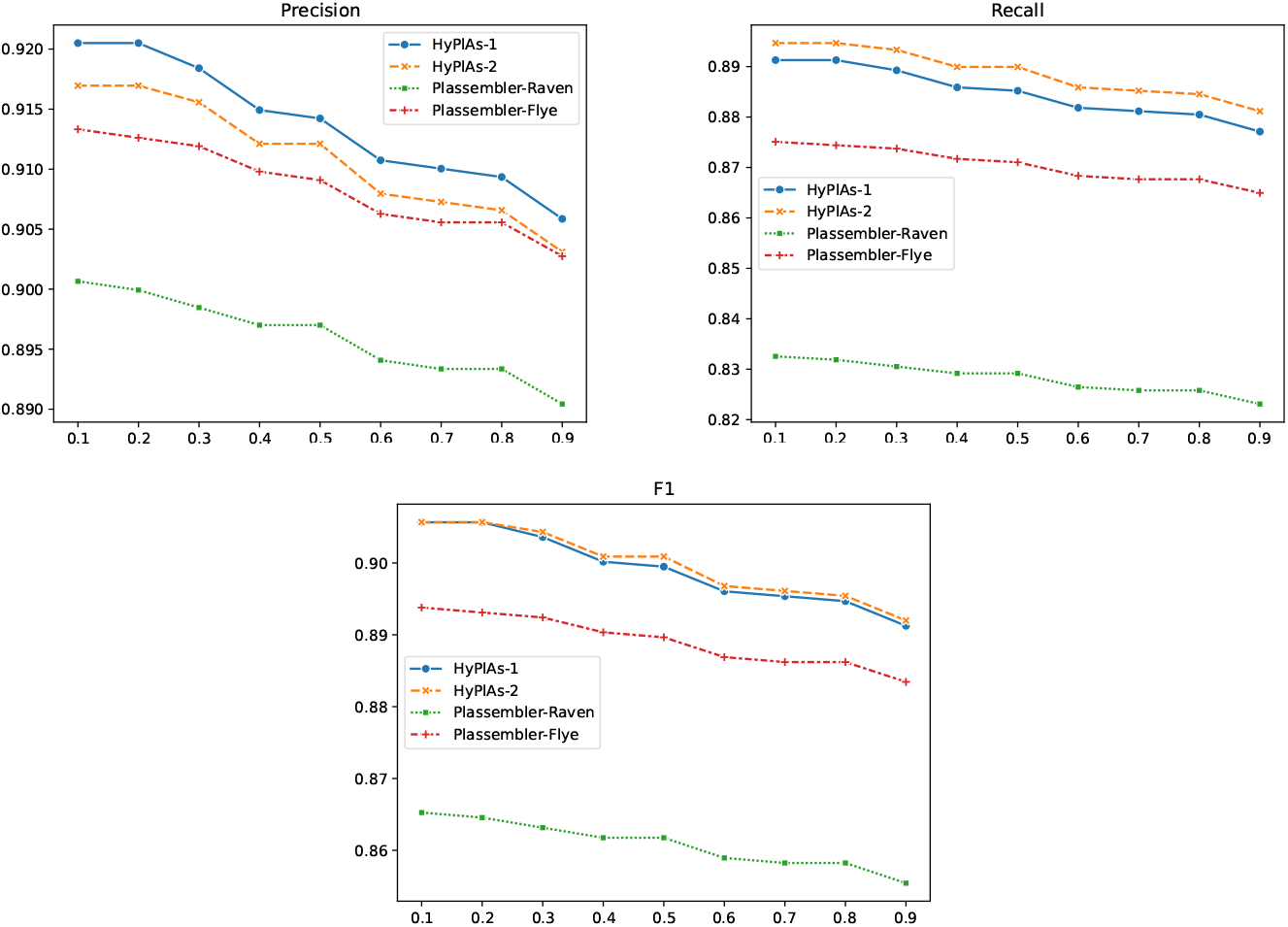
Precision, recall, and F1 scores of Plassembler (with Flye and Raven long-read assemblers) and HyPlAs (with 0, 1 and 2 iterations of the plasmidic long-read augmentation step) over MinHash similarity thresholds (0.1 to 0.9) for a prediction to be accepted as a true positive.

#### Simulated data

For the simulation experiment, we used the reference set for bacterial isolates provided by Bouras et al. [2023] which includes 23 samples including 12 taken from De Maio et al. [2019]. The reference sequences of the 23 bacterial samples were downloaded from Plassembler simulation benchmarking Github repository.

Using these, we simulated long reads using Badread [Wick, 2019] and short reads using a combination of TKSM [Karaoğlanoğlu et al., 2024] and ART [Huang et al., 2012].

Long-read data were generated with the following Badread [Wick, 2019] command:

~~~
badread simulate
  --reference \{input.ref\}
  --small\_plasmid\_bias
  --quantity \{randint(25,35)\}x
~~~

Short-read data were generated with the following commands:

~~~
tksm random-wgs -r \{plasmid.ref\} -o plasmid.mdf --circular --frag-len-dist “normal 350 50” --depth \{randint(30, 2500)\}
tksm random-wgs -r \{chromosome.ref\} -o chromosome.mdf --circular --frag-len-dist “normal 350 50” --depth \{randint(30, 50)\}
cat plasmid.mdf chromsome.mdf $|$
tksm sequence -r {input.ref} -i /dev/stdin --perfect short\_frag.fa -O fasta
art\_illumina -ss HS20 -amp -mp -na -l 100 -f 1 -s 10 -qs 10 -qs2 10 -i short\_frag.fa -o \{output.prefix\}
~~~

Out of 87 simulated plasmids, HyPlAs was able to recover 80 plasmids with 1 false positive while Plassembler (with Flye) recovered 74 (see Table 2). This analysis only covered the complete plasmid assemblies, where incomplete (non-circular) contigs reported by Plassembler are not included in the true or false positive set.

**Table 2.**
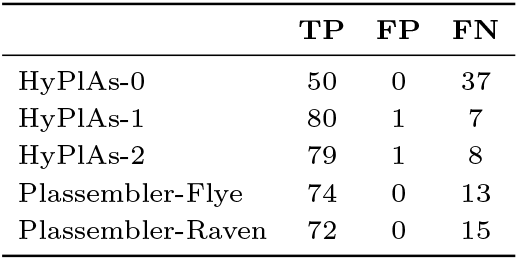
Number of true positives (TP), false positives (FP), and false negatives (FN) on the simulated dataset (87 plasmids).

